# Outbreak of Highly Pathogenic Avian Influenza H5N1 in New England Seals

**DOI:** 10.1101/2022.07.29.501155

**Authors:** Wendy Puryear, Kaitlin Sawatzki, Nichola Hill, Alexa Foss, Jonathon J. Stone, Lynda Doughty, Dominique Walk, Katie Gilbert, Maureen Murray, Elena Cox, Priya Patel, Zak Mertz, Stephanie Ellis, Jennifer Taylor, Deborah Fauquier, Ainsley Smith, Robert A. DiGiovanni, Adriana van de Guchte, Ana Silvia Gonzalez-Reiche, Zain Khalil, Harm van Bakel, Mia K. Torchetti, Julianna B. Lenoch, Kristina Lantz, Jonathan Runstadler

## Abstract

The recent incursion of Highly Pathogenic Avian Influenza A (H5N1) virus into North America and subsequent dissemination of virus across the continent, has had significant adverse impacts on domestic poultry, and has led to widespread mortality in many wild bird species. Here we report the recent spillover of H5N1 into marine mammals in the northeastern United States, with associated mortality on a regional scale. This spillover is coincident with a second wave of H5N1 in sympatric wild birds also experiencing regional mortality events. Viral sequences derived from both seal and avian hosts reveal distinct viral genetic differences between the two waves of infection. Spillover into seals was closely related to virus from the second wave, and one of eight seal-derived sequences had the mammalian adaptation PB2 E627K.

**One-Sentence Summary:** An outbreak of H5N1 in New England seals is the first known population-scale mammalian mortality event associated with the emerging highly pathogenic avian influenza clade 2.3.4.4b.

## Main Text

Few questions in infectious disease research are more critical to public health than identifying how and why pathogens cross species barriers and emerge in mammals. Seals are predominantly colonial marine mammals that share habitat with coastal waterbirds. Harbor (*Phoca vitulina*) and gray (*Halichoerus grypus*) seals in the North Atlantic are known to be affected by avian influenza A virus (IAV) and have experienced prior outbreaks involving seal-to-seal transmission (1–5). These seal species represent a pathway for adaptation of IAV to mammalian hosts that has proven to be a recurring event in nature with implications for human health. Harbor seals have been shown to be particularly susceptible to IAV morbidity and mortality. Gray seal populations have had milder IAV outbreaks than harbor seals, but also have measurable influenza antibodies, even during periods of no observed outbreaks or excess mortality and may represent a reservoir-like host of some influenza subtypes (6, 7).

Highly pathogenic avian influenza (HPAI) viruses are of major concern for their pandemic potential and the socioeconomic impact of agricultural outbreaks. Of special concern are IAV subtypes H5 and H7 due to the potential to mutate to HPAI. Specifically, the goose/Guangdong H5 HPAI viruses, which emerged in 1996, are the only HPAI viruses known to be sustained in wild waterfowl populations (8). Since October 2020, H5N1 HPAI belonging to the goose/Guangdong H5 2.3.4.4b clade has been responsible for over 70 million poultry deaths across Africa, Asia, Europe, and North America (9). As of July 2022, the World Organisation for Animal Health (WOAH) has reported more than 100 wild mammal infections with (H5) clade 2.3.4.4b in many mesocarnivore species including seals and foxes (9–14). Rare human infections with H5 clade 2.3.4.4b viruses have been reported (15–17). To date, there have been no reports of onward transmission of H5 clade 2.3.4.4b in mammalian species. Here we report an outbreak of H5N1 HPAI among New England harbor and gray seals concurrent with a second wave of avian infections in the region that is of a large enough scale to be categorized as a seal Unusual Mortality Event (UME).

The first North American infections with HPAI clade 2.3.4.4b were from samples collected in November 2021 in Canada and late December 2021 in the US (18, 19). Phylogenetic analysis supports at least one incursion of H5 2.3.4.4b via the Atlantic flyway (20, 21). As of July 13, 2022, there have been 126 federally reported wild bird detections in New England (**Fig. 1A, table S1**). Starting on January 1, 2022 avian oropharyngeal and/or cloacal samples were collected for viral surveillance from wild birds through four wildlife rehabilitation facilities in Massachusetts. Additionally, opportunistic samples were collected in Maine and Massachusetts in response to suspicious avian deaths observed on seabird breeding colonies (**Fig. 1A, table S2**). Materials and methods are available as supplementary materials.

**Fig. 1.**
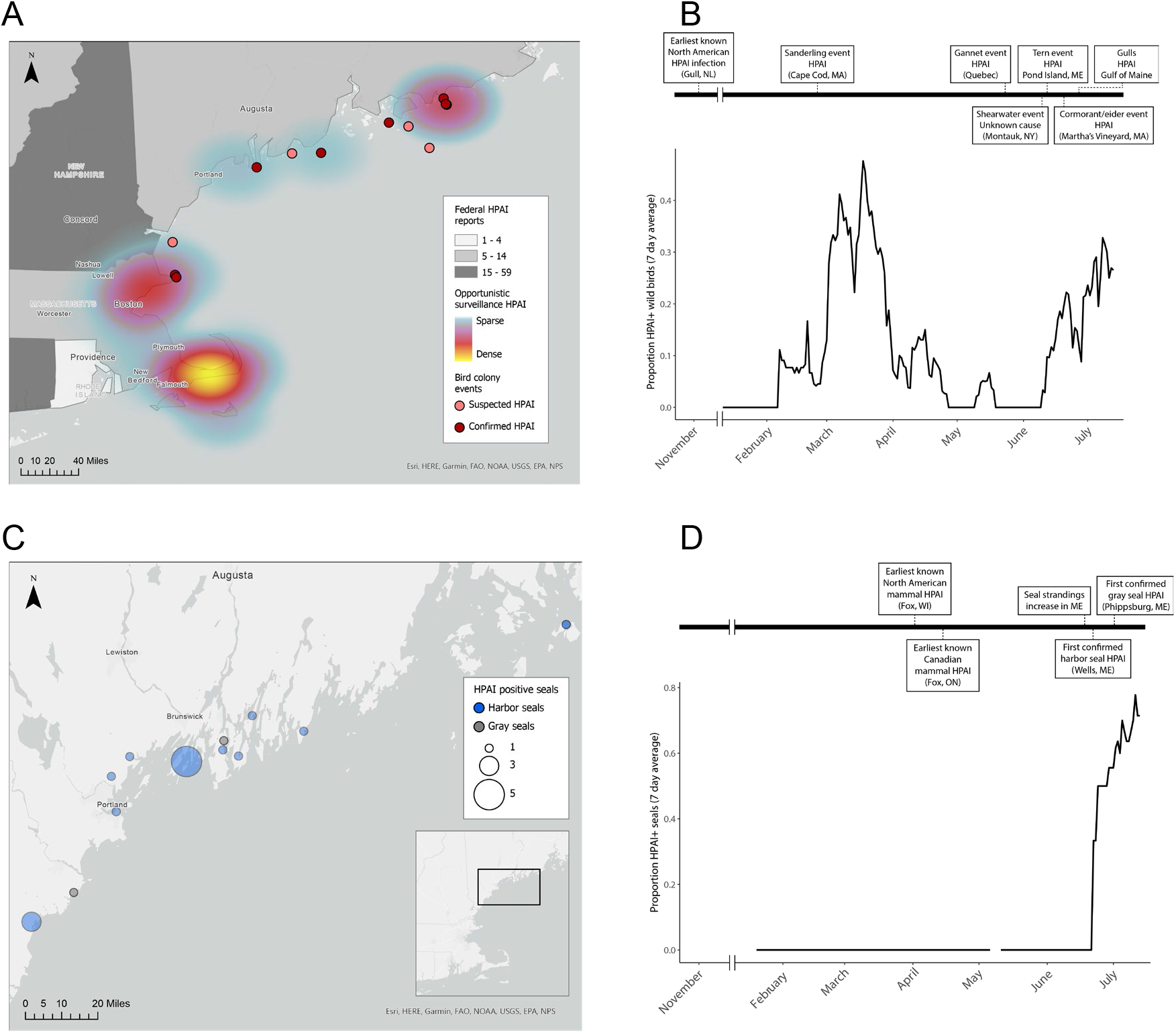
Opportunistic surveillance of New England birds and seals for H5N1 HPAI. Widespread detection of HPAI in New England began in February 2022. (**A**) Geographic distribution of observed HPAI in New England birds. Federal reports of HPAI are shown at the state level, shaded in gray. Regional opportunistic surveillance is shown as a heat map where the bird or sample was collected, and bird colonies with suspected or confirmed HPAI mass mortalities are samples shown as dots. (**B**) Rolling 7-day average of H5 positive birds by RT-PCR from Massachusetts rehabilitation facilities and opportunistic field collection (n=869 unique birds). Two waves of HPAI have been observed in the region to date. A timeline of observed avian mortality events in New England and the Canadian Maritimes during this period illustrates an increased number of population mortality events in the second wave. (**C**) Location of stranded or deceased HPAI positive harbor seals (blue circles) and gray seals (gray circles). Circles are proportional to the number of animals found at that location. (**D**) Rolling 7-day average of H5 positive seals by RT-PCR from US Atlantic coast stranded animals (n=121 unique seals). The first detection of HPAI in wild seals was from samples collected in late June (harbor seal) and early July (gray seal).

We screened samples from 869 individual wild birds representing 73 avian species of concern for H5 influenza and identified 105 infected birds from 19 species (**Fig. 1B, data S1**). Birds screened through the rehabilitation centers had no symptom-specific inclusion criteria, enabling detection of asymptomatic infection and an overview of regional trends. These data show that New England has experienced two waves of infections in wild birds during 2022. The first wave peaked in March and was largely represented by raptor mortalities (37% of positives). A second wave began in June with gulls and eiders being most frequently reported (35.1% and 31.6% of positives, respectively). Additionally, mortality events affecting seabird breeding colonies throughout the coastal region were reported during the second wave, with eight islands having at least one bird test positive for H5.

Concurrent with the second wave of avian infections, increased seal strandings and carcasses found on shore were observed in Maine starting in mid-June. These seal mortalities were declared by NOAA to be an Unusual Mortality Event retrospective to June 1, 2022 (22). Through routine surveillance we screened 121 pinnipeds (67 harbor seals, 34 gray seals, 20 harp seals) between January 20 and July 13, 2022 (**data S1**). Nasal, oral, conjunctival and/or rectal samples were collected along the North Atlantic coast from Maine to Virginia with no symptom-specific inclusion criteria. From January through June 14^th^, there were no detections of HPAI in any of the 92 stranded seals that were tested.

On June 21^st^, a juvenile harbor seal that stranded in Wells, Maine was the first case determined to be positive for HPAI. A juvenile gray seal that stranded in Phippsburg, ME on July 1^st^ was the first of that species positive for HPAI. From June 21^st^ through July 13^th^, a total of 15 of 25 harbor seals and 2 of 4 gray seals with HPAI were detected along the coast of Maine. The HPAI positive seals were within coastal regions of known and suspected HPAI outbreaks among terns, eiders, cormorants and gulls (**Fig. 1C,D**). The majority of stranded seals were deceased. Of those that stranded live, symptoms included respiratory signs with a subset of neurologic cases. The respiratory tract was the most consistent source of RT-PCR positive sample type from affected animals (17/18 nasal, 15/17 oral, 7/17 conjunctiva, 6/18 rectal).

Influenza A viruses were sequenced directly from samples resulting in 55 avian (53 complete, 2 partial), and 8 seal (all complete) viral genomes from New England (Accession: XXXXXX-XXXXXX). Sequences were analyzed using the vSNP pipeline (https://github.com/USDA-VS/vSNP) and RAxML to generate a phylogenetic tree and table of single nucleotide polymorphisms (SNPs). All samples from the second wave of avian infections were genetically distinct from first wave viruses, and seal-derived viruses clustered with the second avian wave with strong support (**Fig. 2A**). During the second avian infection wave, the seal derived viruses clustered closest to eiders, cormorant, gulls and some raptors, all of which had observed mortality in the New England region during this period. Further resolution of the sub-clades associated with each wave were not well supported, as viruses in these groups were highly similar. Therefore, the precise ancestry of the seal-origin viruses should not be overinterpreted. One highly divergent branch composed of virus from first wave raptors was excluded from the vSNP analysis but is shown in **figure S1**. Notably, this group includes three second wave terns, which were part of a mortality event on the breeding grounds on Pond Island, Phippsburg, Maine.

**Fig. 2.**
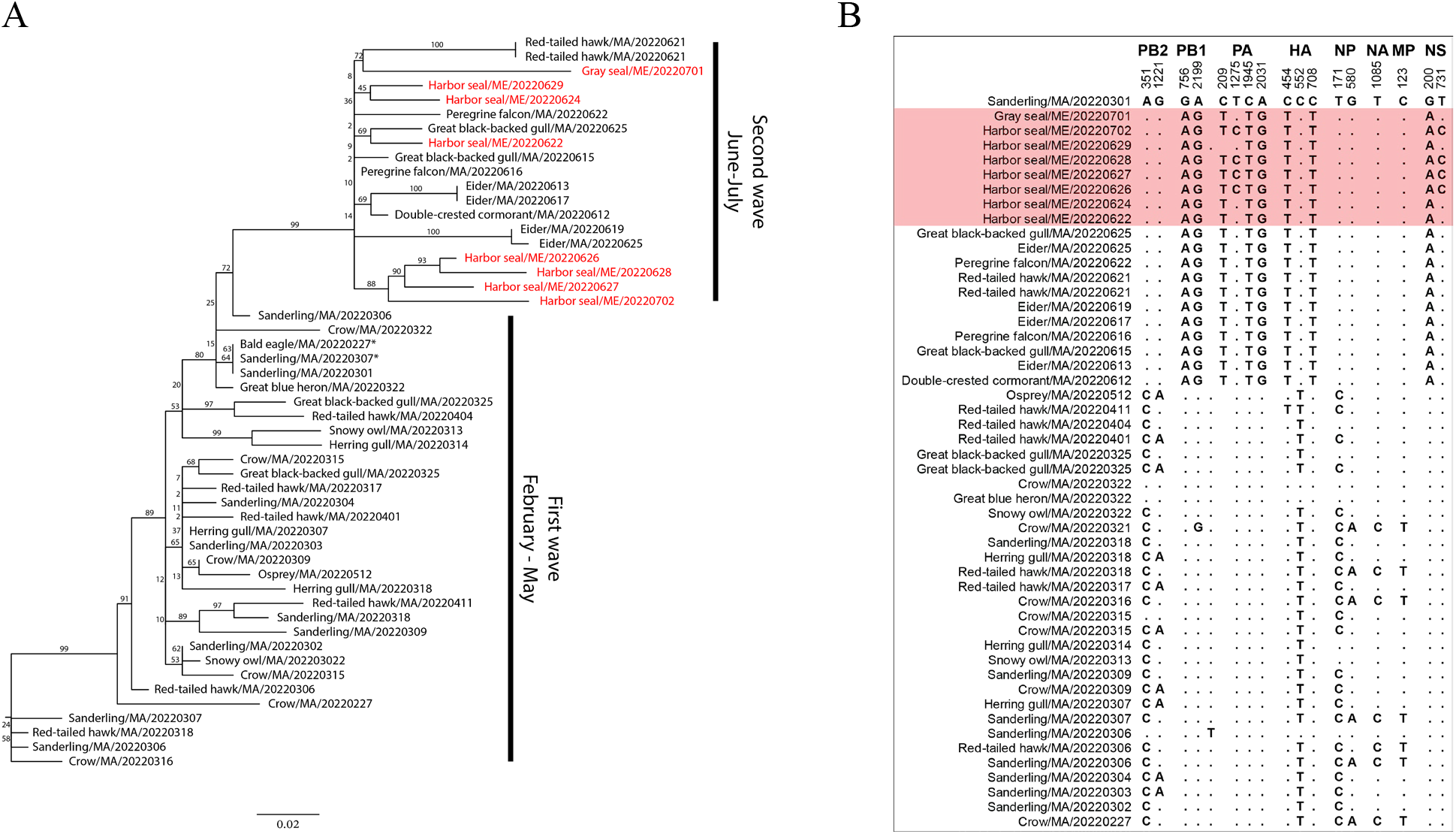
Genetic analysis of New England bird and seal origin H5N1 HPAI. Complete and partial genomes (asterisk) of H5N1 HPAI were compared using the vSNP pipeline with Sanderling/MA/20220301 as a reference. All specimens were collected in the New England region from February to July 2022. (**A**) RAxML was run on vSNP output with bootstrap=10000. Support values are labeled on branches and seal-derived sequences are colored red. (**B**) Comparative SNP output was separated into seals and birds, sorted in reverse chronological order and subset to relevant positions. Seal-derived sequences are highlighted red. Influenza segments and nucleotide position within segments labeled on top. Consensus with Sanderling/MA/20220301 at a position is symbolized with a dot.

Sequences of virus from four harbor seals clustered with high confidence. These seals share two rare SNPs, PA:T1275C and NS:T731C, which have not yet been observed in any New England birds (Fig. 2B). PA:T1275C is a silent mutation and is present in 17/857 (2%) of currently available comparable sequences, including 0/13 mammals. NS:T731C results in amino acid change NS:I244T. This mutation is present in 36/859 (4.2%) of currently available comparable sequences, including 2/12 foxes (**table S3, data S1**). The four seals from which these viruses were detected all originated from different towns and spanned a geographic region of approximately 30 miles. Of the eight seal-derived sequences, one had the PB2 substitution E627K that has been associated with mammalian adaptation. This sequence was obtained from the nasal swab of a deceased juvenile harbor seal in Harpswell, ME on June 28th, 2022, just one week after the first HPAI detection in seals. PB2:E627K has been observed in 2/849 (0.24%) of currently available comparable sequences, and these are 2/12 foxes (**table S4, data S1**). None of the sequences contained previously described PB2 substitution D701N that is also associated with mammalian adaptation (23). Other nucleotide differences observed in multiple seals are consistent with second wave avian viruses.

Since the initial incursion of HPAI H5N1 2.3.4.4b into North America during the last year, the virus has spread south and west across the continent, affecting both domestic and wild avian species, as well as several species of terrestrial mammals, and has now spilled over into marine mammals in the northeastern US. Mammals have generally been considered dead-end hosts for HPAI; however, given the extent of new species infections during the 2022 event and the high proportion of scavenging species affected, further transmission via scavenging or predation is possible. It is currently unclear if the marine mammal spillover will also be a dead-end transmission event, but given the early detection of a sequence with evidence of mammalian adaptation and the extent of mortality already associated with the species, the marine mammal spillover appears more impactful to these species than the sporadic terrestrial mammal spillovers observed to date.

Unlike other documented spillovers of HPAI H5N1 into terrestrial wild mammals, it is unlikely that multiple seals acquired virus through predation or scavenging of an infected source, as birds are not a typical food source for harbor or gray seals (24). Transmission is likely occurring through either environmental transmission or direct contact between seals, though current data is unable to distinguish between these two possible routes. Uniquely different from other seal outbreaks of IAV for which dabbling ducks, including Mallard (*Anas platyrhynchos*) or Blue-winged Teal (*Anas discors*) have been identified as a possible source host, this H5N1 outbreak identifies a novel transmission pathway between marine birds and mammals. The assemblage of host species in the North Atlantic, and the life-history or anthropogenic factors that underlie their susceptibility, may be critical for predicting future outbreaks in seals with implications for human health.

The spillover into seals in 2022 may be linked to behavior and seasonal ecology of birds and seals. Colony-associated bird mortality events have been more observed during the summer/second wave which could create a dense pool of infectious birds and carcasses that overlap with seal haul-out sites. Given the ongoing detections of HPAI in seabirds (gulls, terns) and sea ducks (eider) throughout the New England coast, it is possible that virus shed in the feces of congregated birds may serve as a source of infection for seals via oral, nasal or conjunctival routes (25). If individual bird-seal spillover events represent the primary transmission route, the associated seal Unusual Morality Event suggests that this mode of transmission is occurring frequently and with a low species barrier for seals.

Alternately, seasonal seal behavior, along with extensive exposure to infected birds, may have resulted in one or more seal infections with onward seal transmission. The data herein include viruses from four seals that share two unusual mutations which have not yet been observed in any New England birds. This may be explained by (1) unobserved avian variants in the population, (2) strong host-specific pressure that selects for these mutations in an infected seal, or (3) seal-to-seal transmission following one or more spillover infections. As the seal-specific mutation pattern is only present in half of the seals (4 of 8) and within the same branch, it is unlikely that host pressure is playing a major role.

Migratory waterfowl are natural reservoirs for influenza A viruses, allowing long range and transcontinental movement of these viruses (26). Marine mammals are known to be susceptible to a wide range of influenza subtypes that can cause large scale mortality events, or asymptomatic circulation within the population (6). Harbor seals haul-out in population clusters for pupping season in May through early June, and perhaps missed the first wave of HPAI in New England. However, these seals haul-out in even greater numbers and in dense colonies during molting periods from July-August. During these periods, harbor seals will still travel, generally within 50km but over 100km from their haul-out sites, potentially allowing geographical spread of HPAI among the species (27, 28). In addition, harbor and gray seals share haul-out sites throughout the Gulf of Maine and prior to the gray seal pupping season in early winter, when large numbers of that species congregate together.

Unlike in agricultural settings, outbreaks in wild populations cannot be controlled or managed well through biosecurity measures or depopulation. This is particularly true of large, mobile marine species like seals. Colonial wildlife, avian and mammalian, may be particularly impacted by influenza A viruses and may allow for ongoing circulation between and within species. This provides opportunity for reassortments of novel strains and mammalian adaptation of virus. Migratory animals may then disseminate virus over broad geographic regions. The wild interface of coastal birds and marine mammals is therefore critical for monitoring influenza A viruses of pandemic potential, particularly in light of the impossibility of replicating such an interface in a research environment.

## Acknowledgments

We are grateful to the numerous staff and volunteers of Marine Mammals of Maine, College of the Atlantic, Seacoast Science Center, National Marine Life Center, Marine Mammal Alliance Nantucket, International Fund for Animal Welfare, Atlantic Marine Conservation Society, NY Marine Rescue Center, National Aquarium, Mystic Aquarium, Tufts Wildlife Clinic, New England Wildlife Center, New England Fisheries and Science Center, Wild Care Inc., Cape Wildlife, Linda Loring Nature Foundation, Maine Coastal National Wildlife Reservation, UMass Nantucket Field Station, U.S. Fish and Wildlife Services, and MA Audubon who provide critical expertise on marine and avian wildlife, obtain samples from stranded animals, and provide essential insights on ecological context from the field. The scientific results and conclusions, as well as any views or opinions expressed herein, are those of the authors and do not necessarily reflect the views or policies of the U.S. Government, its agencies, or any of the included organizations.

## Author contributions

Conceptualization: WP, KS

Methodology: WP, KS, AGR

Investigation: WP, KS, AF, LD, DW, KG, MM, EC, PP, ZM, SE, JT, RG

Formal analysis: WP, KS, AG, AGR, ZK, HB, MT, KL

Visualization: KS

Funding acquisition: WP, JR

Project administration: WP, KS, LD, MM, PP, ZM, SE, DF, AS, RG, HB, MT, JR

Supervision: WP, KS, JR

Writing – original draft: WP, KS

Writing – review & editing: WP, KS, NH, AF, JS, LD, DW, KG, MM, EC, PP, ZM, SE, JT, DF, AS, RG, AG, AGR, ZK, HB, MT, JL, KL, JR

## Data and materials availability

All data are available in the main text or the supplementary materials.

## Supplementary Materials

Materials and Methods

Fig. S1

Tables S1 to S4

Data S1

## References and Notes

1. J. R. Geraci et al., Mass mortality of harbor seals: pneumonia associated with influenza A virus. Science 215, 1129–1131 (1982).

2. R. J. Callan, G. Early, H. Kida, V. S. Hinshaw, The appearance of H3 influenza viruses in seals. J Gen Virol 76 (Pt 1), 199–203 (1995).

3. S. J. Anthony et al., Emergence of fatal avian influenza in New England harbor seals. mBio 3, e00166–00112 (2012).

4. R. Bodewes et al., Avian Influenza A(H10N7) Virus-Associated Mass Deaths among Harbor Seals. Emerging Infectious Diseases 21, 720–722 (2015).

5. R. Bodewes et al., Spatiotemporal Analysis of the Genetic Diversity of Seal Influenza A(H10N7) Virus, Northwestern Europe. J Virol 90, 4269–4277 (2016).

6. W. B. Puryear et al., Prevalence of influenza A virus in live-captured North Atlantic gray seals: a possible wild reservoir. Emerg Microbes Infect 5, e81 (2016).

7. R. Bodewes et al., Seroprevalence of Antibodies against Seal Influenza A(H10N7) Virus in Harbor Seals and Gray Seals from the Netherlands. PLoS One 10, e0144899 (2015).

8. R. Harfoot, R. J. Webby, H5 influenza, a global update. J Microbiol 55, 196–203 (2017).

9. World Organisation for Animal Health (2022) World Animal Health Information System.

10. D. L. Shin et al., Highly Pathogenic Avian Influenza A(H5N8) Virus in Gray Seals, Baltic Sea. Emerging Infectious Diseases 25, 2295–2298 (2019).

11. T. Floyd et al., Encephalitis and Death in Wild Mammals at a Rehabilitation Center after Infection with Highly Pathogenic Avian Influenza A(H5N8) Virus, United Kingdom. Emerging Infectious Diseases 27, 2856–2863 (2021).

12. A. Postel et al., Infections with highly pathogenic avian influenza A virus (HPAIV) H5N8 in harbor seals at the German North Sea coast, 2021. Emerg Microbes Infec 11, 725–729 (2022).

13. J. M. Rijks et al., Highly Pathogenic Avian Influenza A(H5N1) Virus in Wild Red Foxes, the Netherlands, 2021. Emerging Infectious Diseases 27, 2960–2962 (2021).

14. A. Schulein, M. Ritzmann, J. Christian, K. Schneider, A. Neubauer-Juric, Exposure of wild boar to Influenza A viruses in Bavaria: Analysis of seroprevalences and antibody subtype specificity before and after the panzootic of highly pathogenic avian influenza viruses A (H5N8). Zoonoses Public Hlth 68, 503–515 (2021).

15. World Health Organization (2022) Human infection with avian influenza A (H5N8) - Russian Federation (World Health Organization).

16. World Health Organization (2022) Influenza A (H5) - United Kingdom of Great Britain and Northern Ireland (World Health Organization).

17. Centers for Disease Control and Prevention (2022) U.S. Case of Human Avian Influenza A(H5) Virus Reported. (Centers for Disease Control and Prevention).

18. Canadian Food Inspection Agency National Emergency Operation Centre Geographic Information System Services (Highly Pathogenic Avian Influenza - Wild Birds. (Canadian Food Inspection Agency National Emergency Operation Centre Geographic Information System Services,).

19. United States Department of Agriculture (USDA Confirms Highly Pathogenic Avian Influenza in a Wild Bird in South Carolina. (United States Department of Agriculture).

20. S. N. Bevins et al., Intercontinental Movement of Highly Pathogenic Avian Influenza A(H5N1) Clade 2.3.4.4 Virus to the United States, 2021. Emerg Infect Dis 28, 1006–1011 (2022).

21. V. Caliendo et al., Transatlantic spread of highly pathogenic avian influenza H5N1 by wild birds from Europe to North America in 2021. Sci Rep 12, 11729 (2022).

22. National Oceanic and Atmospheric Administration (2020) 2018–2020 Pinniped Unusual Mortality Event Along the Northeast Coast (Office of Protected Resources, National Oceanic and Atmospheric Administration).

23. G. Gabriel, V. Czudai-Matwich, H. D. Klenk, Adaptive mutations in the H5N1 polymerase complex. Virus Res 178, 53–62 (2013).

24. W. D. Bowen, G. D. Harrison, Comparison of harbour seal diets in two inshore habitats of Atlantic Canada. Can J Zool 74, 125–135 (1996).

25. G. Martin, D. J. Becker, R. K. Plowright, Environmental Persistence of Influenza H5N1 Is Driven by Temperature and Salinity: Insights From a Bayesian Meta-Analysis. Front Ecol Evol 6 (2018).

26. N. J. Hill et al., Ecological divergence of wild birds drives avian influenza spillover and global spread. PLoS Pathog 18, e1010062 (2022).

27. R. M. J. Pace, Elizabeth; Wood, Stephanie A.; Murray, Kimberly; Waring, Gordon (2019) Trends and Patterns of Seal Abundance at Haul-out Sites in a Gray Seal Recolonization Zone.

28. S. J. Hayes; E. Maze-Foley, K. Rosel, P. Byrd B. Cole T. Henry A (2019) US Atlantic and Gulf of Mexico Marine Mammal Stock Assessments - 2019. ed S. J. Hayes, E. Maze-Foley, K. Rosel P (National Oceanic and Atmospheric Administration).

