## Supplementary material for "Outbreak of Highly Pathogenic Avian Influenza H5N1 in New England Seals": table S1, table S2, table S3, table S4, figure S1

**This PDF file includes:**

Materials and Methods

Fig. S1

Tables S1 to S4

Caption for Data S1

**Other Supplementary Materials for this manuscript include the following:**

Data S1

Materials and Methods

Sample Collection

Oropharyngeal and/or cloacal samples were collected from birds that were brought into rehabilitation for care or were found deceased. All live animals were sampled by experienced personnel within the facility (Tufts Wildlife Clinic, Cape Wildlife Clinic, New England Wildlife Centers, or Wild Care, Inc.) as part of diagnostic care. Deceased animals were either sampled in the field or brought to a wildlife center for sampling.

Oral, conjunctival, nasal, and/or rectal samples were collected from seals by staff at Marine Mammals of Maine (MMoME), Seacoast Science Center (SSC), the National Marine Life Center (NMLC), International Fund for Animal Welfare (IFAW), Mystic Aquarium, and National Aquarium. Samples were obtained from both live and dead pinnipeds (gray, harbor, and harp seals) that stranded along the Eastern Atlantic seaboard spanning Maine to Maryland, from January – July 2022. Samples from stranded animals were collected as diagnostic samples under each organizations Stranding Agreement with NOAA Fisheries Service.

Swab samples were collected using polyester swabs (Puritan Medical Products, Guilford, ME, USA) placed into viral transport media (VTM) comprised of Medium 199, nystatin, gentamicin, benzylpenicillin, streptomycin, sulfamethoxazole, kanamycin sulfate (Sigma-Aldrich, St. Louis, MO) and bovine serum albumin (ThermoFisher Scientific, Waltham, MA, USA). Samples were stored at -80°C or -20°C until processed.

RT-PCR

RNA was extracted from 50µl of VTM per sample using the Mag-Bind Viral DNA/RNA 96 kit (Omega Bio-Tek Inc., Norcross, GA, USA) on a semiautomated KingFisher Purification System robot (ThermoFisher Scientific, Waltham, MA, USA) as described in Puryear et al (6). RNA was screened for the influenza A virus (IAV) Matrix (MP) gene by reverse transcription polymerase chain reaction (RT-PCR) on the StepOnePlus platform (ABI, Beverly, MA). 5μl total RNA was added to qScript XLT One-Step RT-qPCR ToughMix ROX (VWR, Franklin, MA) with forward M F25 (5’-AGATGAGTCTTCTAACCGAGGTCG-3’), reverse 2002 M R124 (5’-TGCAAAAACATCTTCAAGTCTCTG-3’) and probe M P64 (FAM-TCAGGCCCCCTCAAAGCCGA-TAM) oligonucleotides and a one-step real time RT-PCR run was performed as follows: 50°C, 10 minutes; 95°C, 1 minute; (95°C, 3 seconds; 60°C, 30 seconds) for 45 cycles. All plates were run with multiple negative VTM controls and purified A/Puerto Rico/8/1934 H1N1 RNA for a positive control. Any sample with a Ct<40 was further screened for Hemagglutinin H5 using the same protocol described above but with forward H5 1456NA (5’-ACGTATGACTATCCACCATACTCA-3’), forward H5 1456EA (5’-ACGTATGACTACCCGCAGTATTCA-3’), reverse H5 1685 (5’-ACCTCGATGGGCAATGTGTT-3’) and probe H5 1637 (FAM-CATGTCCCTCATATCAAAACCTTCGGAGG-TAM) oligonucleotides. Samples with a Ct<35 were sent to the National Veterinary Services Laboratories for further confirmatory RT-PCR testing.

Sequencing

Whole genome sequencing was performed at the Icahn School of Medicine at Mount Sinai (ISMMS) and National Veterinary Services Laboratory (NVSL). IMSSM: Original sample material from 25 animals was provided. RNA was extracted using QIAmp viral RNA minikit (Qiagen, Hilden, Germany) and amplified using a modified two-step RT-PCR. RT was performed with the ProtoScript II kit (New England Biolabs, Ipswich, MA) using 7µL RNA input with 1µL of the Opti1 primer set incubated at 65°C for 5 minutes to denature the RNA after which 10µL of ProtoScript II Reaction Buffer and 2µL of ProtoScript II Reverse Transcriptase was added and incubated at 25°C for 5 min, 48°C for 30min, and 80°C for 5 min to inactivate the enzyme. PCR was performed on cDNA with a Q5 high-fidelity PCR kit (New England Biolabs, Ipswich, MA) according to the manufacturer’s instructions adjusted to a 25µL volume reaction with a 5µL cDNA input using the Opti1 primer set, consisting of primers Opti1-F1 (5´-GTTACGCGCCAGCAAAAGCAGG), Opti1-F2 (5´-GTTACGCGCCAGCGAAAGCAGG), and Opti1-R1 (5´-GTTACGCGCCAGTAGAAACAAGG). DNA amplicons were purified using an Agencourt AMPure XP 5ml kit (Beckman Coulter, Pasadena, CA) and prepared using the Nextera XT DNA library prep kit (Illumina, San Diego, CA) according to manufacturer protocol, and sequencing was performed on a MiSeq instrument (Illumina) with 2 × 150-base paired end reads. NVSL: Original sample material from 38 animals was PCR amplified and cDNA libraries prepared using the Nextera XT DNA Sample Preparation Kit according to manufacturer instructions (New England Biolabs, Ipswich, MA). Sequencing was performed using Reagent Kit v2 (500-cycles) on the MiSeq platform (Illumina, San Diego, CA).

Phylogenetic analysis

SNP-based phylogenetic analysis was performed using the NVSL vSNP pipeline. Two datasets were run: (1) All 63 genomes using Chicken/NL/FAV-0033/20221221 as a reference and (2) 52 genomes that fall into more closely related clusters using Sanderling/MA/20220301 as a reference. RAxML version 8.2.12 was run on vSNP output alignment of non-consensus nucleotides using a GTRγ model and bootstrap=10000. Trees are organized by increasing node order.

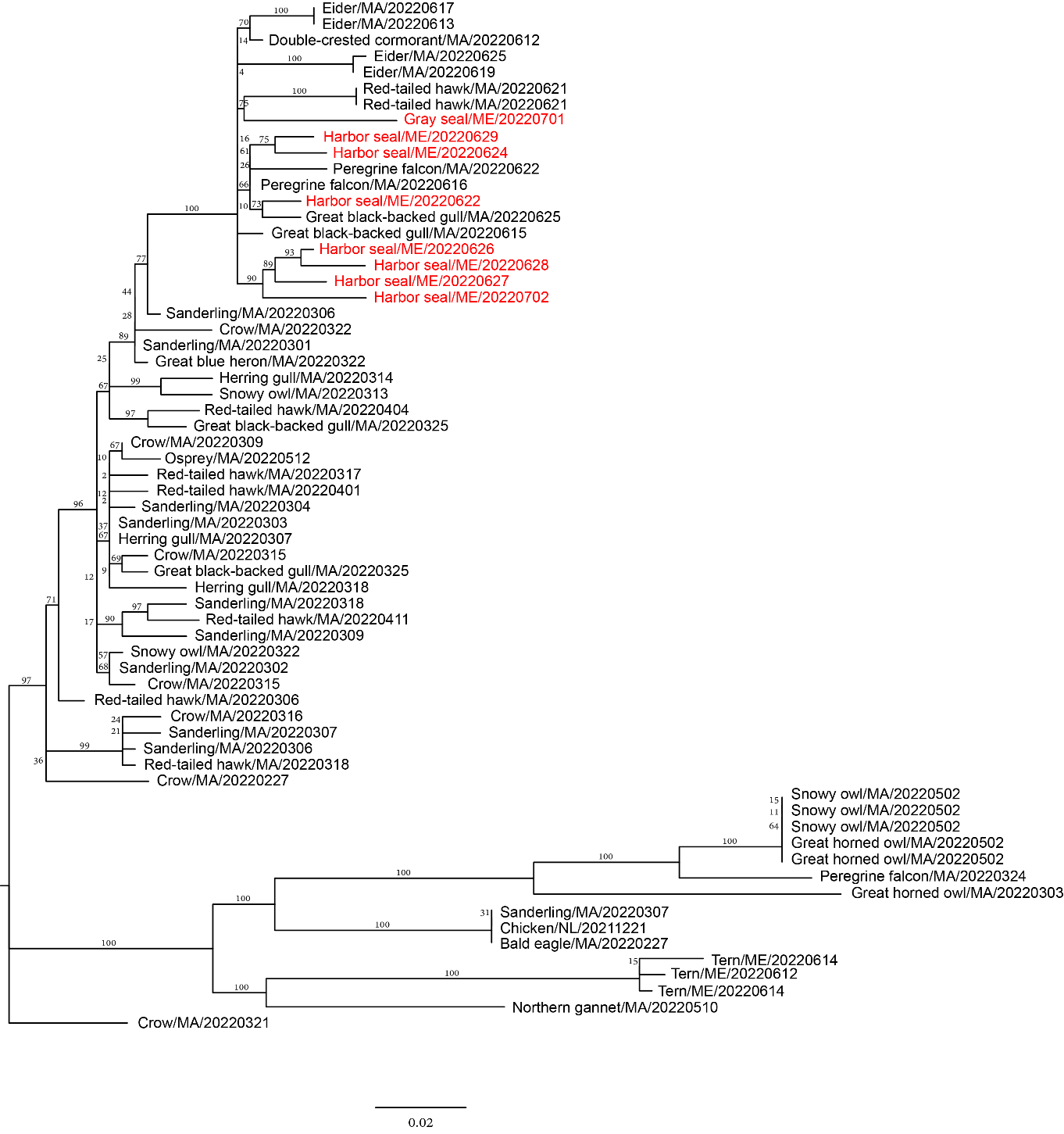

Fig. S1. Genetic analysis of all New England bird and seal origin H5N1 HPAI.
Complete and partial genomes (asterisk) of H5N1 were compared using the vSNP pipeline with Chicken/NL/FAV-0033/20221221 as a reference. All specimens were collected in the New England region from February to July 2022. RAxML was run on vSNP output with bootstrap=10000. Support values are labeled on branches and seal-derived sequences are colored red.

Table S1.

Federally reported cases in wild birds in New England states through July 13, 2022.

| **Collection_date** | **Species** | **State_Province** | **County** |
| --- | --- | --- | --- |
| 2/9/2022 | Mallard | New Hampshire | Rockingham |
| 2/9/2022 | Mallard | New Hampshire | Rockingham |
| 2/9/2022 | Mallard | New Hampshire | Rockingham |
| 2/9/2022 | Mallard | New Hampshire | Rockingham |
| 2/9/2022 | Mallard | New Hampshire | Rockingham |
| 2/9/2022 | Mallard | New Hampshire | Rockingham |
| 2/9/2022 | Mallard | New Hampshire | Rockingham |
| 2/9/2022 | Mallard | New Hampshire | Rockingham |
| 2/9/2022 | Mallard | New Hampshire | Rockingham |
| 2/9/2022 | Mallard | New Hampshire | Rockingham |
| 2/9/2022 | Mallard | New Hampshire | Rockingham |
| 2/9/2022 | Mallard | New Hampshire | Rockingham |
| 2/9/2022 | Mallard | New Hampshire | Rockingham |
| 2/9/2022 | Mallard | New Hampshire | Rockingham |
| 2/9/2022 | Mallard | New Hampshire | Rockingham |
| 2/9/2022 | Mallard | New Hampshire | Rockingham |
| 2/9/2022 | Mallard | New Hampshire | Rockingham |
| 2/9/2022 | Mallard | New Hampshire | Rockingham |
| 2/9/2022 | Mallard | New Hampshire | Rockingham |
| 2/9/2022 | Mallard | New Hampshire | Rockingham |
| 2/16/2022 | Mallard | Connecticut | New London |
| 2/16/2022 | Mallard | Connecticut | Middlesex |
| 2/16/2022 | Mallard | Connecticut | Middlesex |
| 2/16/2022 | Mallard | Connecticut | Middlesex |
| 2/16/2022 | Mallard | Connecticut | Middlesex |
| 2/16/2022 | Mallard | Connecticut | Middlesex |
| 2/16/2022 | Mallard | Connecticut | Middlesex |
| 2/16/2022 | Mallard | Connecticut | Middlesex |
| 2/16/2022 | Mallard | Connecticut | Middlesex |
| 2/16/2022 | Mallard | Connecticut | Middlesex |
| 2/16/2022 | Mallard | Connecticut | Middlesex |
| 2/16/2022 | Mallard | Connecticut | Middlesex |
| 2/16/2022 | Mallard | Connecticut | Middlesex |
| 2/16/2022 | Mallard | Connecticut | Middlesex |
| 2/16/2022 | Mallard | Connecticut | Middlesex |
| 2/16/2022 | Mallard | Connecticut | Middlesex |
| 2/16/2022 | Mallard | Connecticut | Middlesex |
| 2/16/2022 | Mallard | Connecticut | Middlesex |
| 2/16/2022 | Mallard | Connecticut | Middlesex |
| 2/16/2022 | Mallard | Connecticut | Middlesex |
| 2/16/2022 | Mallard | Connecticut | Middlesex |
| 2/16/2022 | Mallard | New Hampshire | Grafton |
| 2/16/2022 | Mallard | New Hampshire | Grafton |
| 2/16/2022 | Mallard | New Hampshire | Grafton |
| 2/16/2022 | Mallard | New Hampshire | Rockingham |
| 2/16/2022 | Mallard | New Hampshire | Rockingham |
| 2/16/2022 | Mallard | New Hampshire | Rockingham |
| 2/16/2022 | Mallard | New Hampshire | Rockingham |
| 2/16/2022 | Mallard | New Hampshire | Rockingham |
| 2/16/2022 | Mallard | New Hampshire | Rockingham |
| 2/16/2022 | Mallard | New Hampshire | Rockingham |
| 2/16/2022 | Mallard | New Hampshire | Rockingham |
| 2/16/2022 | Mallard | New Hampshire | Rockingham |
| 2/16/2022 | Mallard | New Hampshire | Rockingham |
| 2/16/2022 | Mallard | New Hampshire | Rockingham |
| 2/16/2022 | Mallard | New Hampshire | Rockingham |
| 2/16/2022 | Mallard | New Hampshire | Rockingham |
| 2/16/2022 | Mallard | New Hampshire | Rockingham |
| 2/16/2022 | Mallard | New Hampshire | Rockingham |
| 2/16/2022 | Mallard | New Hampshire | Rockingham |
| 2/16/2022 | Mallard | New Hampshire | Rockingham |
| 2/16/2022 | Mallard | New Hampshire | Rockingham |
| 2/16/2022 | Mallard | New Hampshire | Rockingham |
| 2/16/2022 | Mallard | New Hampshire | Rockingham |
| 2/16/2022 | Mallard | New Hampshire | Rockingham |
| 2/16/2022 | Mallard | New Hampshire | Rockingham |
| 2/16/2022 | Mallard | New Hampshire | Rockingham |
| 2/16/2022 | Mallard | New Hampshire | Rockingham |
| 2/16/2022 | Mallard | New Hampshire | Rockingham |
| 2/16/2022 | Mallard | New Hampshire | Rockingham |
| 2/23/2022 | American black duck | Connecticut | New Haven |
| 2/23/2022 | American black duck | Connecticut | New Haven |
| 2/23/2022 | American black duck | Connecticut | New Haven |
| 2/23/2022 | American black duck | Connecticut | New Haven |
| 2/23/2022 | American black duck | Connecticut | New Haven |
| 2/23/2022 | American black duck | Connecticut | New Haven |
| 2/23/2022 | American black duck | Connecticut | New Haven |
| 2/23/2022 | American black duck | Connecticut | New Haven |
| 2/23/2022 | American black duck | Connecticut | New Haven |
| 2/23/2022 | American black duck | Maine | Washington |
| 2/23/2022 | American black duck | Maine | Washington |
| 2/23/2022 | American black duck | Maine | Washington |
| 2/23/2022 | American black duck | Maine | Washington |
| 2/23/2022 | American black duck | Maine | Washington |
| 2/23/2022 | American black duck | Maine | Washington |
| 3/1/2022 | Canada goose | Massachusetts | Barnstable |
| 3/1/2022 | Canada goose | Massachusetts | Barnstable |
| 3/1/2022 | Mallard | New Hampshire | Rockingham |
| 3/2/2022 | Canada goose | New Hampshire | Strafford |
| 3/2/2022 | Canada goose | New Hampshire | Strafford |
| 3/2/2022 | Canada goose | New Hampshire | Strafford |
| 3/9/2022 | American black duck | Connecticut | New London |
| 3/9/2022 | American black duck | Connecticut | New London |
| 3/15/2022 | Canada goose | Massachusetts | Middlesex |
| 3/15/2022 | Snowy owl | New Hampshire | Rockingham |
| 3/16/2022 | Sanderling | Massachusetts | Barnstable |
| 3/16/2022 | Sanderling | Massachusetts | Barnstable |
| 3/16/2022 | Sanderling | Massachusetts | Barnstable |
| 3/16/2022 | Sanderling | Massachusetts | Barnstable |
| 3/16/2022 | Sanderling | Massachusetts | Barnstable |
| 3/16/2022 | Sanderling | Massachusetts | Barnstable |
| 3/16/2022 | Sanderling | Massachusetts | Barnstable |
| 3/16/2022 | Sanderling | Massachusetts | Barnstable |
| 3/16/2022 | Red-tailed hawk | Massachusetts | Barnstable |
| 3/16/2022 | Turkey vulture | Massachusetts | Barnstable |
| 3/24/2022 | Canada goose | Maine | York |
| 3/31/2022 | Bald eagle | Vermont | Chittenden |
| 3/31/2022 | Bald eagle | Vermont | Grand Isle |
| 4/6/2022 | Bald eagle | Maine | Lincoln |
| 4/6/2022 | Bald eagle | Maine | York |
| 4/7/2022 | Bald eagle | Vermont | Franklin |
| 4/14/2022 | Canada goose | New Hampshire | Belknap |
| 4/14/2022 | Canada goose | New Hampshire | Belknap |
| 4/15/2022 | Sanderling | Massachusetts | Barnstable |
| 4/19/2022 | Turkey vulture | Vermont | Washington |
| 4/20/2022 | Bald eagle | New Hampshire | Rockingham |
| 4/26/2022 | Canada goose | Vermont | Washington |
| 5/13/2022 | Wood duck | Vermont | Addison |
| 5/18/2022 | Bald eagle | New Hampshire | Merrimack |
| 5/18/2022 | Bald eagle | New Hampshire | Sullivan |
| 5/23/2022 | Bald eagle | Vermont | Orange |
| 6/3/2022 | Red-tailed hawk | Vermont | Bennington |
| 6/3/2022 | Canada goose | Vermont | Addison |
| 6/3/2022 | Bald eagle | Vermont | Essex |
| 6/3/2022 | Canada goose | Vermont | Washington |
| 7/6/2022 | Great black-backed gull | Rhode Island | Washington |

Table S2.

New England seabird breeding colonies where suspicious avian mortalities were observed.

| **Island** | **Location**  **(Town, State, GPS)** | **Mortalities** | **HPAI** |
| --- | --- | --- | --- |
| Ship Island | Steuben, ME  44.4334, -67.89743 | Eiders | Confirmed |
| Green Island | Steuben, ME  44.37341, -67.87305 | Gulls, Eiders, Cormorants | Confirmed |
| Petit Manan Island | Steuben, ME  44.3673, -67.86527 | Gulls, Eiders | Confirmed |
| Swans Island | Swans Island, ME  44.18795, -68.44547 | Gulls | Confirmed |
| Great Duck Island | Frenchboro, ME  44.155, -68.24985 | Gulls | Suspected |
| Mount Desert Rock | Mt Desert Rock, ME  43.96869, -68.12778 | Gulls | Suspected |
| Metinic Island | Vinalhaven, ME  43.88708, -69.12581 | Gulls, terns | Confirmed |
| Pond Island NWR | Phippsburg, ME  43.73952, -69.77064 | Terns | Confirmed |
| Appledore Island | Appledore Island, ME  42.98678, -70.61424 | Gulls | Suspected |
| Thacher Island NWR | Rockport, MA  42.63904, -70.57491 | Gulls | Confirmed |
| Straitsmouth Island | Rockport, MA  42.65981, -70.59115 | Eiders | Confirmed |

Table S3.

Sequences available through July 13, 2022 with an observed T731C substitution in NS. The additional available sequences (N=823) preserve T731 (Data S1).

A/chicken/Italy/IZSLT-122448_21VIR9218-1/2021|2021-10-28|EPI_ISL_7733644

A/chicken/South_Dakota/22-008704-001-original/2022|2022-03-22|EPI_ISL_11971486

A/chicken/South_Dakota/22-008704-002-original/2022|2022-03-22|EPI_ISL_11971487

A/cignus_olor/Italy/IZSLT_21VIR10529-1/2021|2021-12-01|EPI_ISL_8882212

A/Cygnus_olor/Romania/16381_21VIR10306/2021|2021-11-11|EPI_ISL_8440175

A/duck/Bulgaria/756-4_22VIR778-6/2021|2021-12-01|EPI_ISL_11007540

A/duck/Poland/H188_22VIR2515-2/2022|2022-03-02|EPI_ISL_11922808

A/egret/France/21P013418/2021|2021-12-03|EPI_ISL_8377254

A/fox/Iowa/22-014421-001-original/2022|2022-05-06|EPI_ISL_13052721

A/fox/Minnesota/22-014182-001-original/2022|2022-04-22|EPI_ISL_13052720

A/goose/Italy/IZSLT-21VIR10273/2021|2021-11-22|EPI_ISL_7733645

A/great_egret/Czech_Republic/23609/2021|2021-11-28|EPI_ISL_8515483

A/grey_heron/Czech_Republic/23608/2021|2021-11-28|EPI_ISL_8515481

A/grey_heron/Czech_Republic/23608-1K/2021|2021-11-28|EPI_ISL_8515482

A/grey_heron/Czech_Republic/25338-1/2021|2021-12-18|EPI_ISL_12223688

A/grey_heron/Czech_Republic/25338-2/2021|2021-12-18|EPI_ISL_12223734

A/hen/Bulgaria/722-1_22VIR778-1/2021|2021-11-15|EPI_ISL_11007538

A/hen/Bulgaria/757-6_22VIR778-7/2021|2021-12-02|EPI_ISL_11007541

A/hen/Bulgaria/854-1_22VIR778-10/2021|2021-12-29|EPI_ISL_11007537

A/laying_hen/Moldova/68-1_22VIR638-1/2022|2022-01-03|EPI_ISL_11007527

A/laying_hen/Moldova/68-2_22VIR638-2/2022|2022-01-03|EPI_ISL_11007721

A/laying_hen/Romania/10470_22VIR2749-5/2022|2022-02-10|EPI_ISL_11922819

A/mallard/Italy/21VIR6957-6/2021|2021-08-20|EPI_ISL_7987333

A/mallard/New_York/22-008760-007-original/2022|2022-03-22|EPI_ISL_11971491

A/partridge/Bulgaria/745_22VIR778-3/2021|2021-11-23|EPI_ISL_11007722

A/pheasant/New_York/22-008760-008-original/2022|2022-03-22|EPI_ISL_11971490

A/pheasant/New_York/22-009066-001-original/2022|2022-03-25|EPI_ISL_11971502

A/swan/Poland/MB078_22VIR2515-7/2022|2022-02-10|EPI_ISL_11922813

A/swan/Romania/16905_22VIR2749-1/2021|2021-12-08|EPI_ISL_11922815

A/turkey/Bulgaria/755-1_22VIR778-4/2021|2021-11-30|EPI_ISL_11007539

A/turkey/Minnesota/22-010085-001-original/2022|2022-04-03|EPI_ISL_13285555

A/turkey/Minnesota/22-010085-002-original/2022|2022-04-03|EPI_ISL_13285556

A/turkey/Minnesota/22-010312-001-original/2022|2022-04-04|EPI_ISL_13433352

A/turkey/Minnesota/22-010312-004-original/2022|2022-04-04|EPI_ISL_13433365

A/turkey/South_Dakota/22-008485-001-original/2022|2022-03-21|EPI_ISL_11971481

A/turkey/South_Dakota/22-008485-002-original/2022|2022-03-21|EPI_ISL_11971482

Table S4.

Sequences available through July 13, 2022 with an observed E627K amino acid substitution in PB2. The additional available sequences (N=847) preserve E627 (Data S1).

A/fox/Ireland/3866_22VIR2064-1/2022|2022-02-14|EPI_ISL_11259283

A/Fox/Netherlands/EMC1/2022|2022-01-26|EPI_ISL_12066188

Data S1. (separate file)

(1) Table of avian and seal samples screened for influenza A virus and H5 and (2) complete GISAID dataset used for amino acid and SNP comparison with attributions.
